# Maternal Hepatocytes Heterogeneously and Dynamically Exhibit Developmental Phenotypes Partially via YAP1 during Pregnancy

**DOI:** 10.1101/2022.07.27.501729

**Authors:** Shashank Manohar Nambiar, Joonyong Lee, Huaizhou Jiang, Guoli Dai

## Abstract

**Background and Aims:** Pregnancy induces reprogramming of maternal physiology to support fetal development and growth. Maternal hepatocytes undergo hypertrophy and hyperplasia to drive maternal liver growth and alter their gene expression profiles simultaneously. This study aimed to further understand maternal hepatocyte adaptation to pregnancy.

**Methods:** Timed pregnancies were generated in mice.

**Results:** In a non-pregnant state, most hepatocytes expressed *Cd133, α-fetalprotein* (*Afp*,) and *epithelial cell adhesion molecule* (*Epcam*) mRNAs, whereas overall, at the protein level, they exhibited a CD133^-^/AFP^-^ phenotype; however, pericentral hepatocytes were EpCAM^+^. As pregnancy advanced, while most maternal hepatocytes retained *Cd133, Afp*, and *Epcam* mRNA expression, they generally displayed a phenotype of CD133^+^/AFP^+^, and EpCAM protein expression was switched from pericentral to periportal maternal hepatocytes. In addition, we found that the Hippo/yes-associated protein (YAP) pathway does not respond to pregnancy. *Yap1* gene deletion specifically in maternal hepatocytes did not affect maternal liver growth or metabolic zonation. However, the absence of *Yap1* gene eliminated CD133 protein expression without interfering with *Cd133* transcript expression in maternal livers.

**Conclusions:** We demonstrated that maternal hepatocytes acquire heterogeneous and dynamic developmental phenotypes, resembling fetal hepatocytes, partially via YAP1 through a post-transcriptional mechanism. Moreover, maternal liver is a new source of AFP. In addition, maternal liver grows and maintains its metabolic zonation independent of the Hippo/YAP1 pathway. Our findings revealed a novel and gestation-dependent phenotypic plasticity in adult hepatocytes.

**Synopsis:** We found that maternal hepatocytes exhibit developmental phenotypes in a temporal and spatial manner, similarly to fetal hepatocytes. They acquire this new property partially via yes-associated protein 1.

## Introduction

As pregnancy progresses, different maternal non-reproductive organs undergo various organ-dependent adaptive changes to meet the increasing needs of the developing and growing fetus. In pregnant women, adaptive changes in the maternal brain (1), pancreas (2–5), and spleen (6) have been revealed. Certain regions of the maternal brain undergo considerable structural changes, including a decrease in gray matter volume (1). β-cell number in the maternal pancreas increases significantly throughout gestation (3, 5). Conceivably owing to the increase in maternal blood volume (7, 8), the maternal spleen displays a robust enlargement (6). In rodents, structural and/or functional adaptations to pregnancy have been observed in the maternal brain (9), liver (10–12), pancreas (13–15), heart (16, 17), and spleen (18). This phenomenal reprogramming of maternal physiology is believed to be essential for successfully establishing and maintaining gestation. However, these phenomena remain to be comprehensively characterized.

In 1944, Kennaway et al. first observed that pregnancy induced maternal liver enlargement in mice (19). In the last decades, studies have shown that the maternal liver exhibits two major adjustments to pregnancy. The first is dramatic and gestation stage-dependent changes in gene expression profiles, most prominently on nutrient metabolism, organ development, and cell proliferation (10, 12). The second is maternal hepatocyte hypertrophy and hyperplasia, which occur during the second half of pregnancy (10–12, 20). Our previous study demonstrated that the maternal liver coordinates adaptations of the maternal compartment with the placental compartment to ensure the health of the offspring (21). However, many more questions remain regarding the adjustment of the maternal liver to accommodate fetal development and growth. Therefore, this study aimed to understand further the response of the maternal liver to pregnancy. We found that maternal hepatocytes exhibited phenotypes similar to fetal hepatocytes, partially regulated by yes-associated protein 1 (YAP1).

## Results

### Maternal hepatocytes display heterogeneous and dynamic developmental phenotypes

Liver progenitor cells or fetal hepatocytes express CD133 (22), α-fetal protein (AFP) (23–27), and epithelial cell adhesion molecule (EpCAM) (28–32) during development (33). We previously showed that maternal livers highly express *Cd133* mRNA in pregnant rats (10). This finding prompted us to examine the phenotypes of maternal hepatocytes by using these three markers throughout pregnancy in mice.

Maternal livers upregulated *Cd133* mRNA and protein expression as the pregnancy progressed (Fig. 1A-B). *Cd133* transcript levels increased during the first half of gestation, peaked around mid-gestation, then gradually decreased, and eventually returned to the pre-pregnancy state before parturition (Fig. 1A). Maternal liver CD133 protein was rapidly detected after copulation (gestation day 1), progressively enriched, and abundant during the second half of pregnancy (Fig. 1B). Moreover, in the non-pregnant state, *Cd133* mRNA was detected in almost all hepatocytes and mainly localized in the nucleus (Fig. 1C). During pregnancy (gestation day 15), maternal hepatocytes were *Cd133* transcript-positive in the nucleus and cytoplasm, resembling fetal hepatocytes (Fig. 1C). This mRNA molecule was not detected in the non-pregnant hearts (negative control organ) (Fig. 1C). Before pregnancy, a few hepatocytes around the portal triads expressed CD133 (Fig. 1D). In contrast, during pregnancy, almost all maternal hepatocytes were CD133-positive, similar to the fetal hepatocytes (Fig. 1D). Our data revealed that, in response to pregnancy, maternal hepatocytes increased *Cd133* mRNA expression and exported *Cd133* mRNA from the nucleus to the cytoplasm, exhibiting a CD133 protein-positive phenotype.

**Figure 1:**
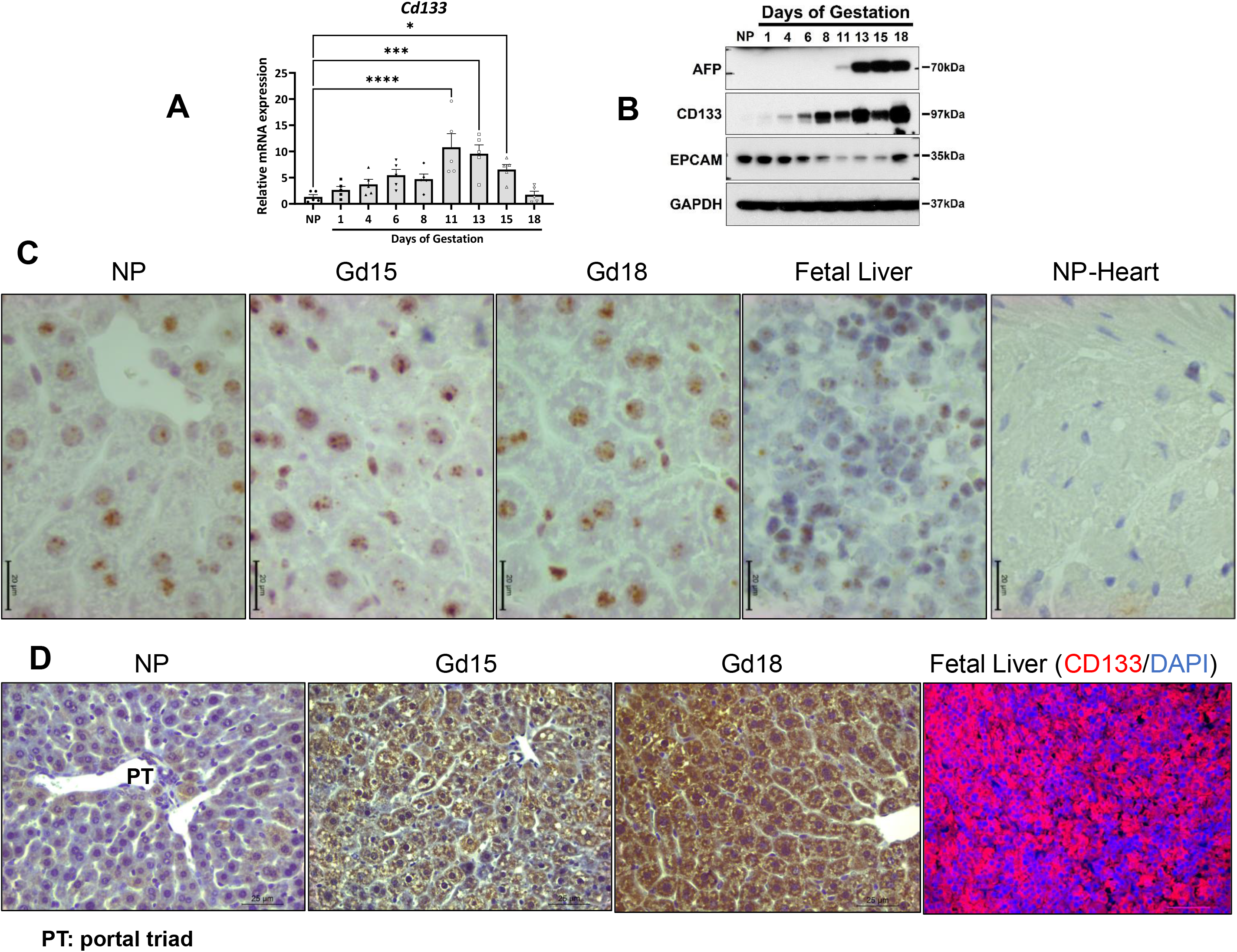
*Cd133* mRNA and protein expression in non-pregnant (NP) and pregnant mouse livers. **(A)** qRT-PCR analysis of maternal hepatic *Cd133* mRNA expression throughout pregnancy. The data indicate the mean fold change relative to NP livers ± SEM (n = 5). *, *P* < 0.05; **, *P* < 0.01 compared to NP livers. **(B)** Western blotting analysis of maternal hepatic CD133, AFP, and EpCAM protein expression throughout gestation. GAPDH was the internal loading control. **(C)** *Cd133* mRNA in situ hybridization of NP and gestation day (Gd) 13 and 15 maternal livers. Fetal livers were a positive control. NP heart was a negative control. *Cd133* mRNA is indicated by brown-colored spots. **(D)** CD133 immunostaining of NP, maternal, and fetal livers.

Relative to the non-pregnant state, maternal livers expressed either unchanged or reduced *Afp* mRNA as pregnancy advanced (Fig. 2A). Inconsistently, maternal livers expressed abundant AFP from mid-gestation to term (Fig. 1B). *Afp* mRNA was detected only in the nuclei of most hepatocytes before pregnancy (Fig. 2B), whereas *Afp* transcripts were expressed in the nuclei and cytoplasm during pregnancy (Fig. 2B). As expected, the non-pregnant heart (negative control organ) was *Afp* mRNA-negative (Fig. 2B), and the AFP protein was undetected in non-pregnant hepatocytes (Fig. 2C). However, AFP protein expression was observed in most maternal hepatocytes after mid-gestation (Gd15) and was subsequently restricted to a subpopulation of maternal hepatocytes, but at an overtly higher expression level (Gd18) (Fig. 2C). In addition, most fetal hepatocytes were positive for *Afp* mRNA in their nuclei and cytosol (Fig. 2B) and positive for AFP protein (Fig. 2C). Together, our data demonstrate that maternal hepatocytes exhibit a phenotype of AFP protein^+^, mimicking fetal hepatocytes, possibly by the exportation of *Afp* mRNA from the nucleus to the cytosol.

**Figure 2:**
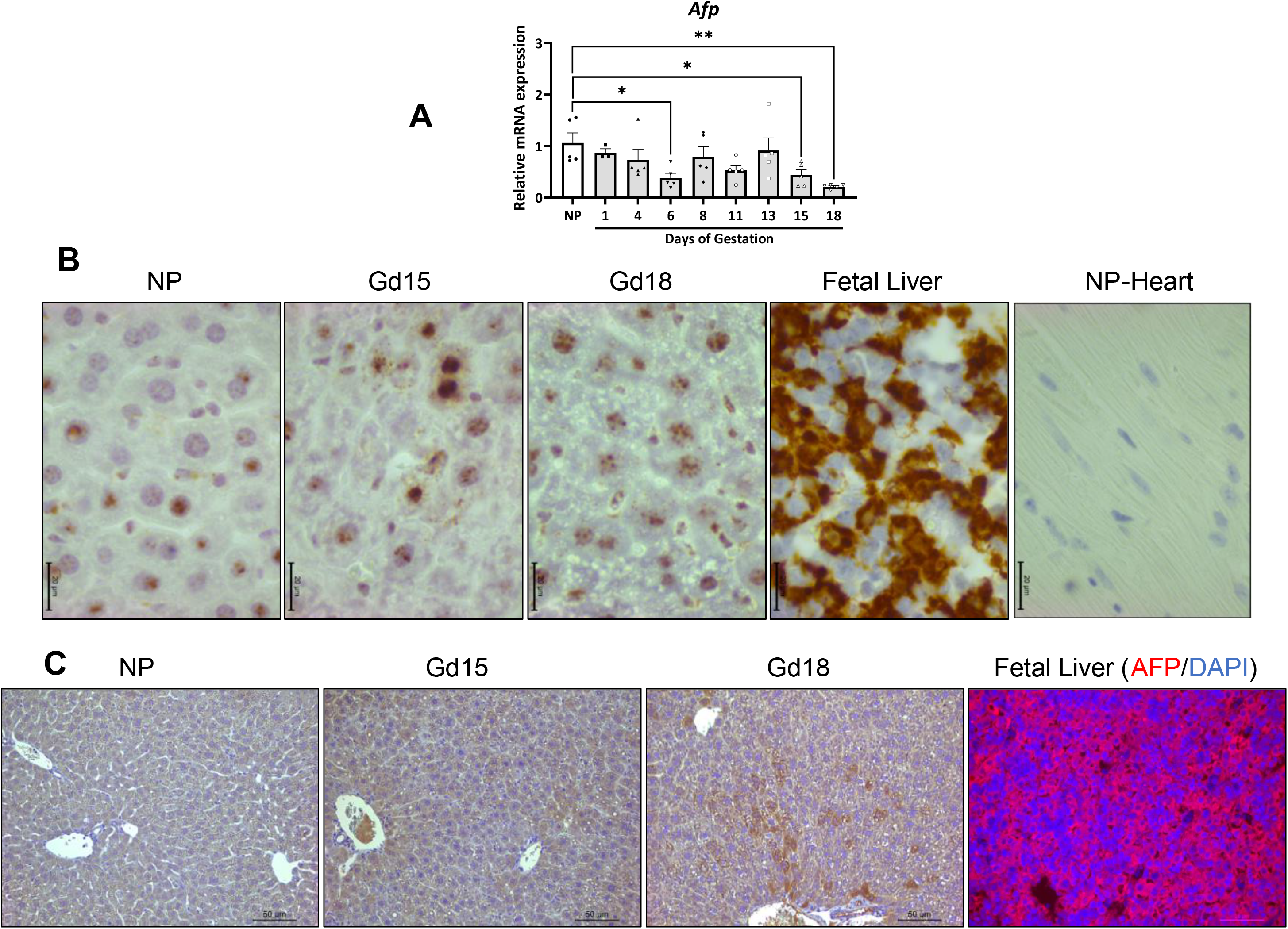
*Afp* mRNA and protein expression in non-pregnant (NP) and pregnant livers. **(A)** qRT-PCR assay of maternal hepatic *Afp* mRNA expression throughout gestation. The data indicate the mean fold change relative to the NP state ± SEM (n = 5). *, *P* < 0.05; **, *P* < 0.01 compared to NP livers. **(B)** *Afp* mRNA in situ hybridization in NP livers and gestation day (Gd) 13 and 15 maternal livers. *Afp* mRNA is indicated by a brown color. Fetal livers were a positive control. NP heart was a negative control. **(C)** AFP immunostaining of NP, pregnant maternal, and fetal livers.

Next, we analyzed *Epcam* mRNA and protein levels in the same setting. Compared with the pre-pregnancy state, *Epcam* mRNA expression was only transiently increased on gestation day 11 in maternal livers (Fig. 3A). However, relative to the non-pregnant state, EpCAM protein expression was overtly reduced after gestation day 6 but returned to equivalent levels before parturition (gestation day 18) (Fig. 1B). In non-pregnant and pregnant livers, *Epcam* mRNA was detected in cholangiocytes, which was expected, and in most hepatocytes, concentrating in the nuclei (Fig. 3B). Regardless of the physiological state, cholangiocytes were EpCAM protein-positive, as anticipated (Fig. 3C). In the non-pregnant state, pericentral hepatocytes were EpCAM protein-positive, whereas periportal hepatocytes were EpCAM protein-negative (Fig. 3C). In striking contrast, during pregnancy, pericentral hepatocytes turned to be EpCAM protein-negative, whereas periportal hepatocytes became EpCAM protein-positive (Fig. 3C). It is known that, during the later liver development stage, EpCAM expression is diminished in fetal hepatocytes and restricted to fetal cholangiocytes (34). Thus, as expected, in embryonic day 18 fetal livers, Epcam mRNA and protein were only detected in cholangiocytes but not in hepatocytes (Fig. 3B-C). Combining these data, we uncovered that pregnancy induces a phenotypic switch from pericentral to periportal maternal hepatocytes, manifested by EpCAM protein expression.

**Figure 3:**
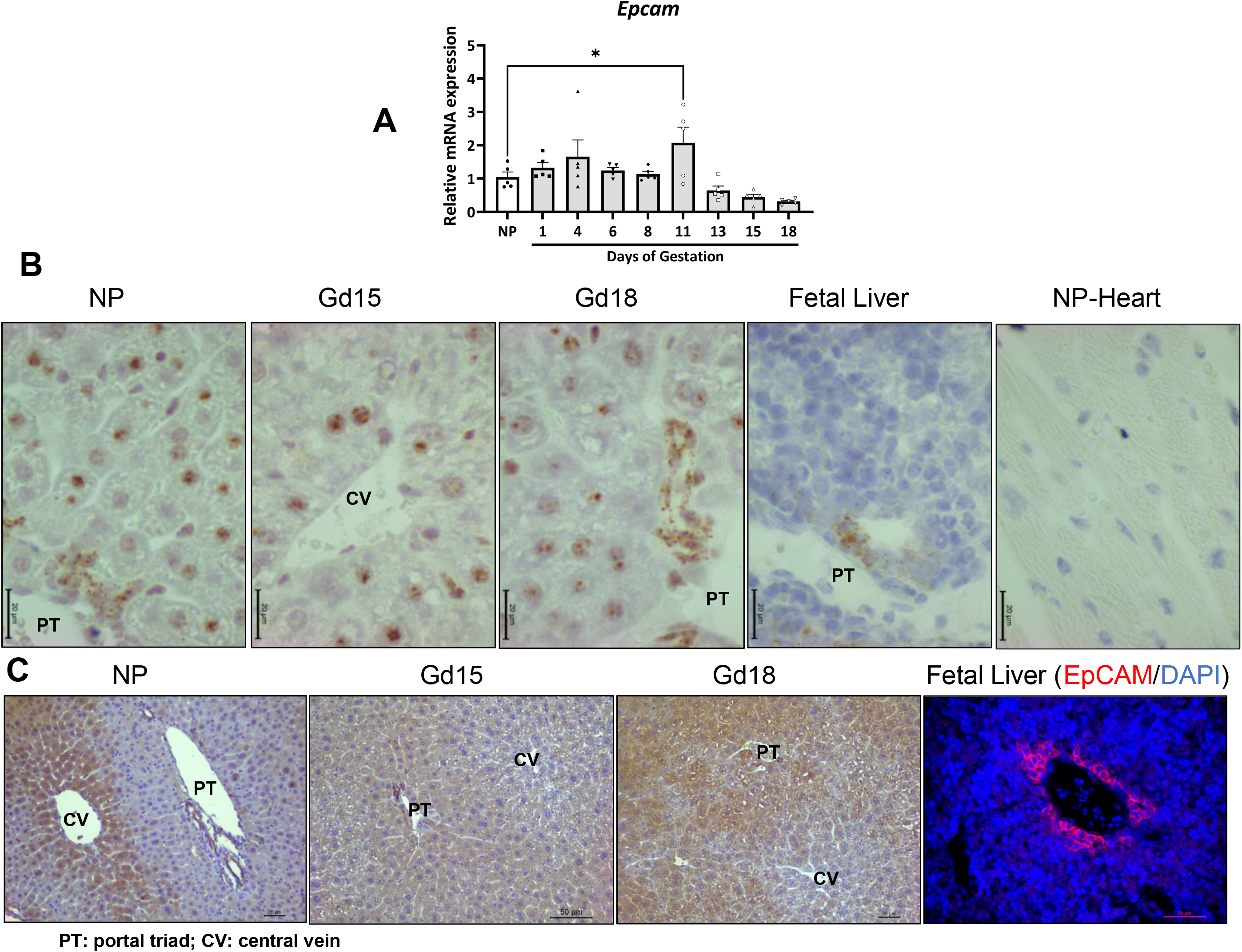
*Epcam* mRNA and protein expression in non-pregnant (NP) and pregnant livers. **(A)** qRT-PCR analysis of maternal hepatic *Epcam* mRNA expression at various stages of gestation. The data indicate the mean fold change relative to the NP state ± SEM (n = 5). *, *P* < 0.05; **, *P* < 0.01 compared to NP livers. **(B)** *Epcam* mRNA in situ hybridization in NP and pregnant maternal livers. *Epcam* mRNA was detected as brown-colored spots. Fetal livers were a positive control. NP heart was a negative control. **(C)** EpCAM immunostaining of NP, pregnant maternal, and fetal livers.

### Pregnancy does not affect the activity of the maternal hepatic Hippo/YAP1 pathway

The canonical Hippo pathway centrally comprises two kinases: MST1/2 and LATS1/2. Phosphorylated and activated MST1/2 phosphorylates and activates LATS1/2. Activated LATS1/2 sequentially phosphorylates and inactivates the transcriptional co-activator YAP or its homolog tafazzin (TAZ), sequestering YAP/TAZ in the cytoplasm. However, dephosphorylation of MST1/2 and subsequent LATS1/2 leads to the dephosphorylation and activation of YAP/TAZ. Active YAP/TAZ is then transported to the nucleus, trans-activating its target genes, including connective tissue growth factor (*Ctgf*) and *Notch2* (35). The Hippo/YAP pathway regulates the growth of organs, including the liver (36), and modulates the phenotypic plasticity of mature hepatocytes (35). Thus, we reasoned that this pathway might control gestation-dependent maternal liver growth and maternal hepatocyte phenotypes. To test this hypothesis, we first evaluated how the components of this pathway respond to pregnancy.

Compared with the non-pregnant state, the expression of maternal hepatic cytoplasmic p-MST1, MST1, p-LAST1, LAST1, p-YAP1, YAP1, p-TAZ, and TAZ was either unchanged or reduced. In addition, the ratios of p-MAST1/total MAST1, p-LAST1/total LAST1, p-YAP1/total YAP1, and p-TAZ/total TAZ were not altered throughout gestation (Fig. 4). Moreover, while the expression of maternal hepatic nuclear YAP1 and TAZ remained the same as before pregnancy (Fig. 5A), the mRNA expression of maternal hepatic *Ctgf* and *Notch2* was unaffected or even reduced as the pregnancy progressed (Fig. 5B). These data indicated that the activity of the Hippo/YAP1 pathway did not show a pregnancy-dependent change.

**Figure 4:**
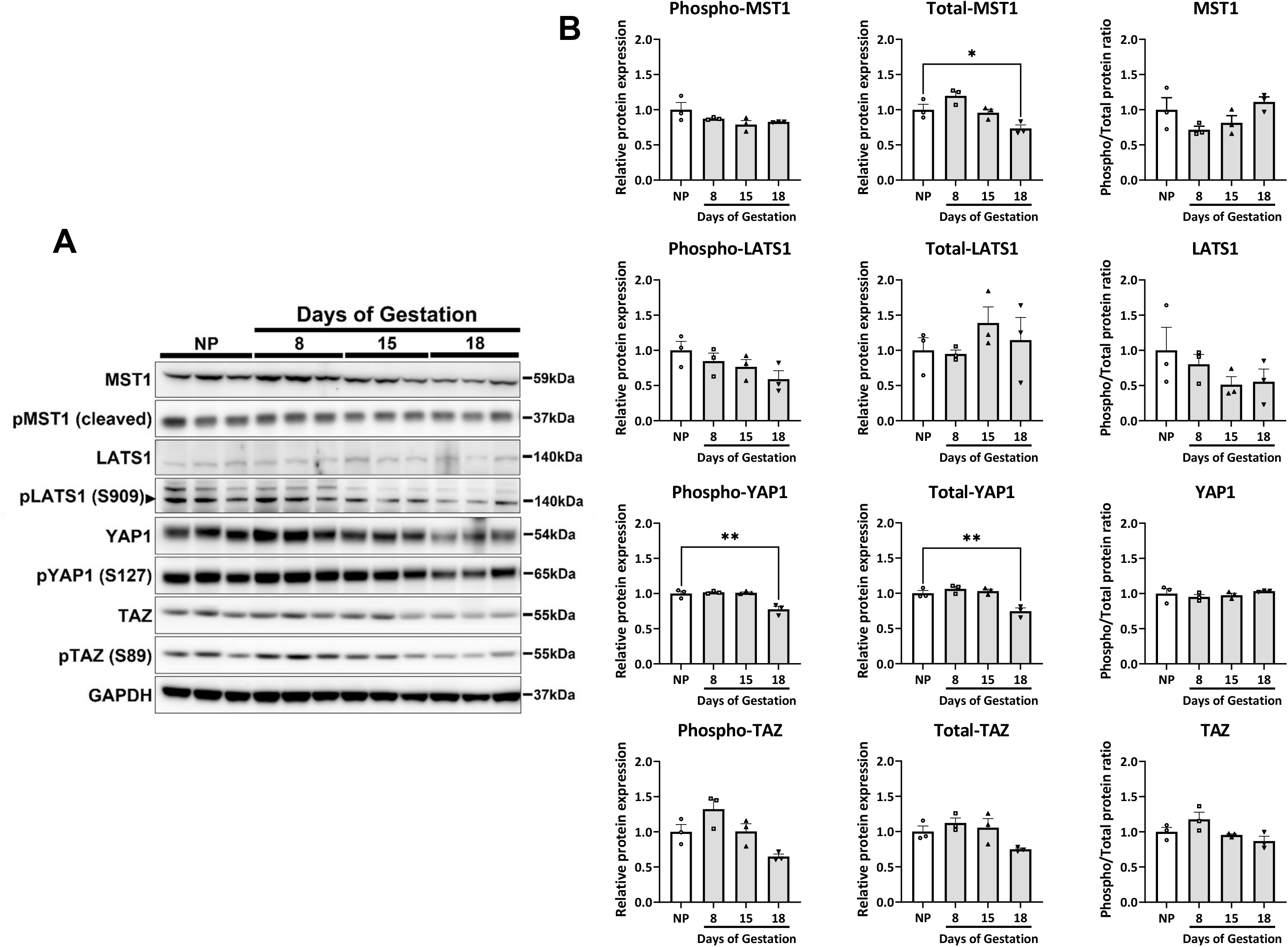
The expression of cytoplasmic components of the Hippo/YAP1 pathway in maternal livers. Timed pregnancies were generated in 3–3.5 months old virgin female C57BL/6J mice. Non-pregnant (NP) and pregnant maternal livers (gestation days 11, 15, and 18) were collected, and cytoplasmic fractions were prepared. **(A)** Western blotting analysis of the cytoplasmic components of the Hippo/YAP pathway. GAPDH was the internal loading control. **(B)** Quantification of relative protein expression of cytoplasmic components of the Hippo/YAP1 pathway. Relative fold changes were calculated after normalizing to GAPDH. Data are indicated as mean ± SEM (n = 3). *, *P* < 0.05 compared to NP livers.

**Figure 5:**
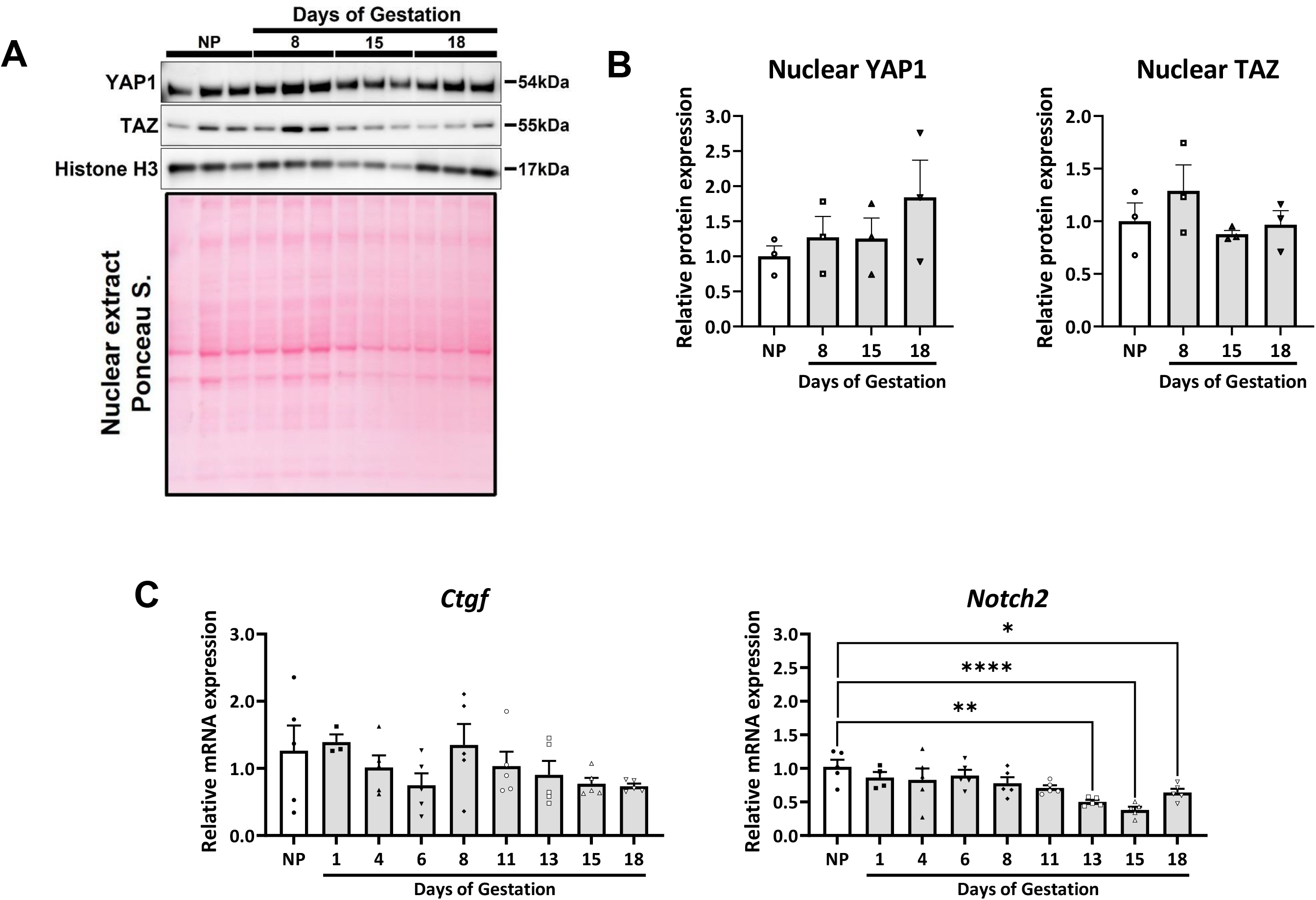
The expression of nuclear YAP1 and TAZ in maternal livers. Timed pregnancies were generated in virgin female C57BL/6J mice. Non-pregnant (NP) and pregnant maternal livers (gestation days 11, 15, and 18) were collected, and nuclear fractions were prepared. **(A)** Western blotting analysis of the nuclear YAP1 and TAZ in maternal livers. The total nuclear protein was stained using ponceau S stain and was the internal loading control. **(B)** Quantification of relative protein expression of nuclear YAP1 and TAZ. Relative fold change was calculated after normalizing to total nuclear protein. Data are shown as mean ± S.E.M (n = 3). **(C)** mRNA expression of maternal hepatic *Ctgf* and *Notch2*. Total mRNA was extracted from NP and pregnant maternal livers (gestation days 1, 4, 6, 8, 11, 13, 15, and 18) of C57BL/6J mice. qRT-qPCR was performed to quantify the mRNA levels of *Ctgf* and *Notch2*. Data represent the mean fold changes relative to NP ± SEM (n = 5). *, *P* < 0.05; **, *P* < 0.01; ***, *P* < 0.001 compared to NP livers.

### The presence of YAP1 is essential for CD133 protein expression in maternal livers

Although pregnancy does not activate maternal hepatic YAP1, we evaluated whether pregnancy-dependent events in the maternal liver require the presence of YAP1. Timed pregnancies were generated in *Yap1*^f/f^ mice by mating them with wild-type male mice. We injected the AAV8-TBG-Cre virus into gestation day 6 *Yap1*^f/f^ mice and deleted the *Yap1* gene specifically in maternal hepatocytes during the second half of pregnancy. The AAV8-TBG-null virus was used as a control. We collected samples on gestation day 18 for various endpoint analyses.

As anticipated, maternal hepatic *Yap1* mRNA and protein expression was largely diminished in maternal hepatocyte-specific *Yap1* knockout (h*Yap1*^-/-^) pregnant mice compared to *Yap1*^f/f^ pregnant ice. The data indicated the high efficiency of the virus-mediated loss of function of *Yap1* (Fig. 6A-D). In the maternal livers without *Yap1*, Cd133 mRNA levels were unaffected, whereas CD133 protein expression almost vanished (Fig. 6A-D). The absence of *Yap1* led to increased *Afp* transcript levels (Fig 6B) but unchanged expression and distribution of AFP protein. The presence or absence of *Yap1* did not affect the expression of *Epcam* mRNA and protein or the distribution of EpCAM protein (Fig. 6A-D). Collectively, these data demonstrate that the lack of *Yap1* post-transcriptionally prevents CD133 protein expression in maternal hepatocytes; therefore, *Yap1* contributes to the regulation of pregnancy-dependent phenotypes of these cells.

**Figure 6:**
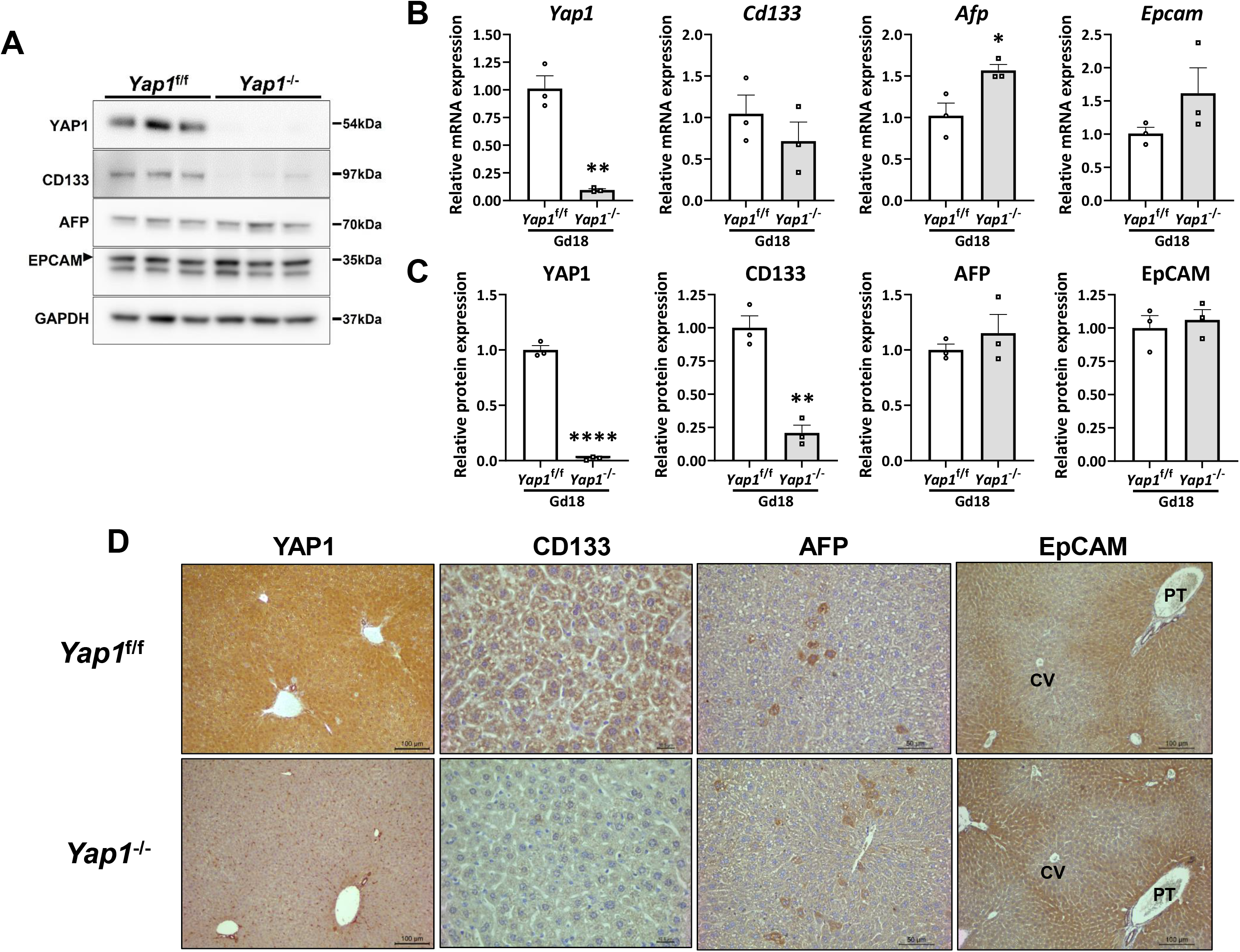
The effects of maternal hepatocyte-specific *Yap1* gene deletion on maternal hepatocyte phenotypes. Timed pregnancies were generated in virgin female *Yap1*^f/f^ mice by mating with wild-type males. On gestation day (Gd) 7, mice were administered with the AAV8-TBG-Cre virus to delete the *Yap1* gene specifically in maternal hepatocytes (*Yap1*^-/-^ mice) or administered the AAV8-TBG-Null virus to serve as the control mice (*Yap1*^f/f^ mice). Gd18 maternal livers were collected from both genotype groups of mice. **(A)** Western blotting analyses of YAP1, CD133, AFP, and EpCAM expression in maternal livers. GAPDH was an internal loading control. **(B)** qRT-PCR analyses of *Yap1, Cd133, Afp, and Epcam* mRNA expression in maternal livers. Data represent mean fold change relative to *Yap1*^f/f^ controls ± SEM (n = 3). *, *P* < 0.05. **(C)** Quantification of YAP1, CD133, AFP, and EpCAM protein expression in maternal livers. Data were normalized with GDPDH. Data represent mean fold change relative to *Yap1*^f/f^ controls ± SEM (n = 3). *, *P* < 0.05; **, *P* < 0.01. **(D)** Immunohistochemistry analyses of YAP1, CD133, AFP, and EpCAM protein distribution in maternal livers.

Furthermore, loss of function of maternal hepatic *Yap1* did not change maternal liver weight, maternal liver-to-total body weight ratio, or maternal hepatocyte density (Fig. 7A). These results implied that *Yap1* might not mediate pregnancy-induced maternal liver growth. Blood chemistry analyses revealed that h*Yap1*^-/-^ pregnant mice displayed increased serum alkaline phosphatase (ALP), decreased serum cholesterol, and unchanged other parameters compared with *Yap1*^f/f^ pregnant mice (Fig. 7B). These data suggested that maternal hepatic *Yap1* is involved in cholesterol metabolism during pregnancy.

**Figure 7:**
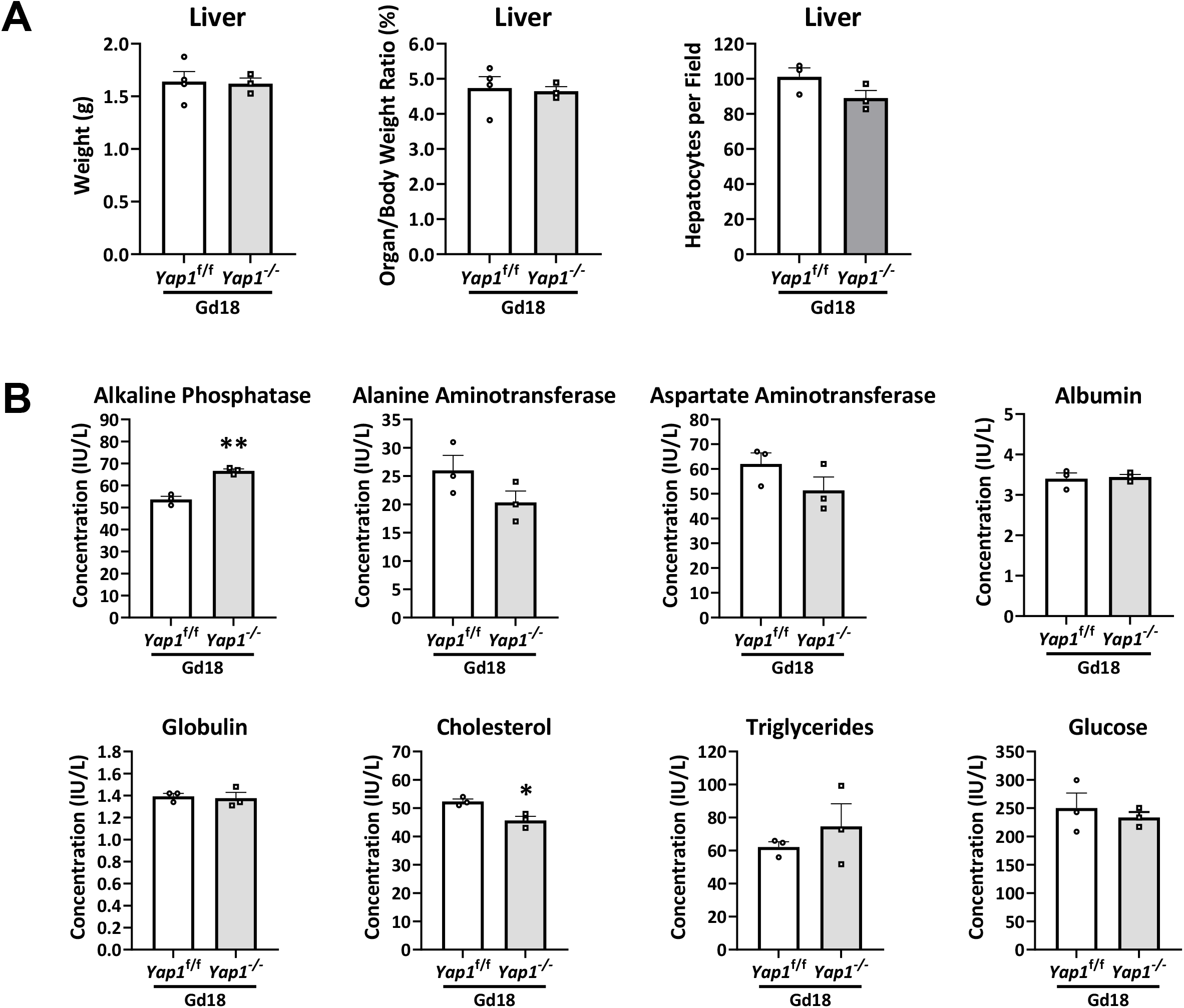
Effect of maternal hepatocyte-specific *Yap1* gene deletion on maternal liver growth and blood chemistry. Timed pregnancies were generated in virgin female *Yap1*^f/f^ mice by mating with wild-type males. On gestation day (Gd) 7, one group of mice was administered the AAV8-TBG-Cre virus to delete the *Yap1* gene specifically in maternal hepatocytes (*Yap1*^-/-^ mice), while the other group was administered the AAV8-TBG-Null virus to act as the control group (*Yap1*^f/f^ mice). Gd18 maternal livers and blood were collected. **(A)** Maternal liver weights, maternal liver-to-total body weight ratios, and maternal hepatocyte densities were shown. **(B)** Maternal serum concentrations of a subset of blood chemistry parameters. Data are shown as mean ± SEM (n = 3). *, *P* < 0.05.

In addition, YAP1 has been shown to regulate liver metabolic zonation (37, 38). Deleting YAP1 from hepatocytes increases zone 3 hepatocytes expressing glutamine synthetase (GS) (38). Therefore, we first evaluated whether pregnancy affects the metabolic phenotypes of maternal hepatocytes by examining the distribution of pericentral hepatocyte markers CYP1A2, CYP2E1, and GS and the periportal hepatocyte markers arginase 1 and E-cadherin. The distribution of these marker proteins did not show overt gestation-dependent changes compared to those in the non-pregnant state (Fig. 8A). We then compared the distribution of these proteins between gestation day 18 *Yap1*^f/f^ and h*Yap1*^-/-^ maternal livers. We also did not observe overt *Yap1*-dependent alterations (Fig. 8B). Hence, these data suggest that neither pregnancy nor YAP1 modulates maternal liver metabolic zonation or hepatocyte metabolic phenotypes.

**Figure 8:**
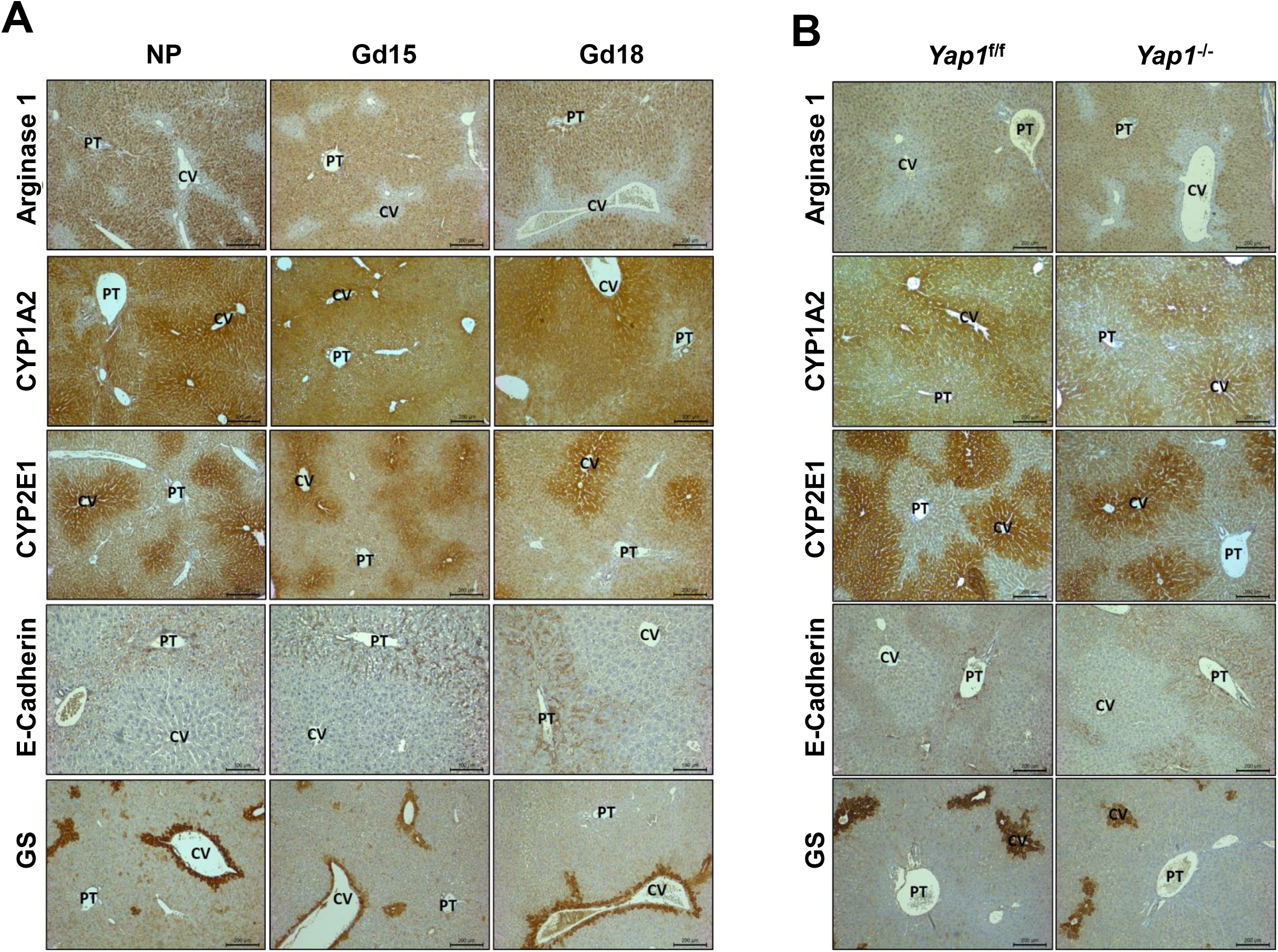
Effect of pregnancy and maternal hepatocyte-specific *Yap1* gene deletion on maternal liver metabolic zonation. Livers of non-pregnant (NP) C57BL/6J mice and maternal livers of gestation day (Gd) 15 and 18 mice of the same strain were collected. **(A)** Distribution of pericentral hepatocyte marker CYP1A2, CYP2E1, and glutamine synthetase (GS) and periportal hepatocyte marker E-cadherin and arginase 1 was analyzed with immunohistochemistry. Timed pregnancies were generated in virgin female *Yap1*^f/f^ mice by mating with wild-type males. On gestation day (Gd) 7, mice were injected with the AAV8-TBG-Cre virus to delete the *Yap1* gene exclusively in maternal hepatocytes (*Yap1*^-/-^ mice) or administered the AAV8-TBG-Null virus to act as the controls (*Yap1*^f/f^ mice). Gd18 maternal livers were harvested from both genotype groups of mice. **(B)** Distribution of CYP1A2, CYP2E1, GS, E-cadherin, and arginase 1 in maternal livers was visualized via immunohistochemistry.

## Discussion

Most interestingly, we discovered for the first time that maternal hepatocytes acquire heterogeneous and dynamic developmental phenotypes during pregnancy. During development, fetal hepatocytes are initially *Cd133*^+^, *Afp*^+^, and *Epcam*^+^ and subsequently *Cd133*^+^, *Afp*^+^, and *Epcam*^-^ at the mRNA level. Correspondingly, fetal hepatocytes display an early-stage CD133^+^/AFP^+^/EpCAM^+^ phenotype and a later-stage CD133^+^/AFP^+^/EpCAM^-^ phenotype at the protein level (25–27, 29, 34). Surprisingly, in the non-pregnant state, most adult female hepatocytes retain the mRNA expression of *Cd133, Afp, and Epcam*. However, in this state, all hepatocytes are AFP^-^, the vast majority of which are CD133^-^, and pericentral hepatocytes are EpCAM^+^. Remarkably, pregnancy induces most maternal hepatocytes to express CD133 and AFP proteins, thus allowing them to exhibit a developmental phenotype of CD133^+^/AFP^+^, simultaneously switching EpCAM protein expression from pericentral to periportal hepatocytes. This process is dynamic. On gestation day 15, most maternal hepatocytes are CD133^+^, whereas, on gestation day 18, all maternal hepatocytes are CD133^+^. In contrast, on gestation day 15, most hepatocytes are AFP^+^, whereas, on gestation day 18, AFP expression is restricted to a randomly distributed subpopulation of maternal hepatocytes. It is well known that adult hepatocytes exhibit heterogeneous metabolic phenotypes. In this study, we revealed heterogeneous developmental phenotypes in this particular physiological state (pregnancy). The exhibition of behaviors by maternal hepatocytes that partially resemble fetal hepatocytes is intriguing. The biological significance of this new feature of adult hepatocytes is unclear and requires further investigation.

Moreover, these pregnancy-induced maternal hepatocyte phenotypic alterations are mediated, at least in part, by the nuclear export of *Cd133* and *Afp* mRNAs. Nuclear retention and export of RNA are highly regulated via various mechanisms and are associated with development, cell fate determination, and cellular functions (39, 40). Before pregnancy, *Cd133* and *Afp* mRNAs were retained in the nuclei of hepatocytes without corresponding protein translation. During pregnancy, they are exported to the cytoplasm of maternal hepatocytes with robust protein production. However, we did not observe nuclear-cytoplasmic export of *Epcam* mRNA in maternal hepatocytes that were positive for EpCAM protein before and during pregnancy. This might be because Epcam transcripts adopt a diffused distribution in the cytosol in hepatocytes; thus, a regular *in situ* hybridization approach is not sensitive enough to detect them. Pregnancy induces a phenotypic switch from pericentral to periportal hepatocytes by inhibiting and allowing EpCAM protein expression in maternal livers, probably via a zone-dependent mechanism.

We also revealed for the first time that maternal hepatocytes produce AFP. AFP is a fetal plasma glycoprotein primarily synthesized by developing fetal hepatocytes and, to a lesser extent, by yolk sac cells. Its presence in the maternal serum is due to fetal and maternal blood exchange during gestation. Clinically, it is used to assess the health of the developing fetus and pregnancy. Abnormally high levels of AFP in maternal serum are considered a potential indicator of various fetal defects or diseases (41). Our findings revealed the maternal liver as an additional source of AFP, implying that abnormally high AFP levels could also be a consequence of maternal liver pathology. Therefore, it is critical to verify this finding in humans. However, obtaining human maternal liver samples is challenging because biopsying this organ is not a routine clinical practice.

The Hippa/YAP pathway modulates liver growth, liver metabolic zonation, and hepatocyte phenotype in adults (42–46). However, our studies indicated that the maternal liver does not rely on this pathway to grow and maintain its metabolic landscape. We arrived at this conclusion based on two pieces of evidence. First, the pathway remains silent in response to pregnancy. Second, deletion of the *Yap1* gene, specifically in maternal hepatocytes, does not affect maternal liver growth or metabolic zonation. However, genetic manipulation of the *Yap1* gene substantially diminished CD133 protein expression without affecting *Cd133* mRNA expression in maternal hepatocytes. Therefore, we can conclude that the presence of YAP1 is indispensable for post-transcriptional expression of CD133 protein and that pregnancy modulates maternal hepatocyte phenotypes partially via YAP1 at the post-transcriptional level. This finding enabled us to gain some mechanistic insight into the regulation of pregnancy-dependent phenotypic plasticity in maternal hepatocytes. In addition, YAP1 has been reported to regulate lipid metabolism, and activation of YAP1 in hepatocytes causes fatty liver (47–50). In this study, we observed that the absence of YAP1 in maternal hepatocytes decreased cholesterol concentrations in the maternal circulation, consistent with these studies. Our findings warrant further studies on how YAP1 is required to maintain pregnant-dependent maternal lipid homeostasis.

In summary, we have demonstrated a novel aspect of pregnancy physiology. As pregnancy progresses, maternal hepatocytes heterogeneously and dynamically adopt developmental phenotypes that mimic fetal hepatocytes, partially via YAP1. In addition, the growth of the maternal liver and maintenance of its metabolic zonation is independent of the Hippo/YAP1 pathway. These findings enabled us to understand further how the maternal liver adapts to pregnancy.

## Materials and Methods

### Animal care and use

Protocol for the care and use of animals was prepared per National Institute of Health guidelines, and all animals were maintained in accordance with the regulations outlined by the Indiana University-Purdue University Indianapolis Animal Care and Use Committee. Animals had ad libitum access to food and drinking water and were fed regular chow. The temperature and relative humidity of the animal rooms were maintained at 22 ± 1°C and 40–60%, respectively. The lighting period comprised 12 h light/dark cycles.

To generate a timed pregnancy, virgin C57BL/6J female and male mice (Jackson Laboratories, Maine, stock no: 000664), aged between 3 and 3.5 months old, were housed together. The following day, the female mouse was inspected for a seminal plug, which indicated successful mating. The appearance of the seminal plug was denoted as gestation day (Gd) 1, and the male was separated from the cage. The *Yap1*^flox/flox^ mouse line was purchased from Jackson Laboratories (stock no: 027929). In this strain, the *Yap1* gene is replaced by the YAP1-flox transgene, and all mice carry this transgene under homozygous conditions. The YAP1-flox transgene is constructed by inserting two loxP sites on either side of the region containing exons-1 and 2 of the YAP1 coding sequence. Maternal hepatocyte-specific YAP1 deletion was achieved by injecting mice with the adeno-associated virus serotype 8 (AAV8) with the thyroxine-binding globulin promoter (*TBG*) promoter expressing *Cre* (AAV8-TBG-Cre virus, Addgene, AV-8-PV1091) via tail vein at a dose of 1×10^12^ genomic copies/mouse. AAV8-TBG-Null virus (Addgene, AV-8-PV0148) was used as a control.

### Quantitative real-time polymerase chain reaction (qRT-PCR)

Total mRNA was extracted from livers (70–100 mg /liver) using TRIzol reagent. Briefly, liver tissues were homogenized using the TRIzol reagent. cDNAs were prepared using a Verso cDNA kit (#AB-1453B, Thermo Scientific) following the manufacturer’s instructions. qRT-PCR was performed using the TaqMan gene expression assay protocol. The master mix solution was prepared using molecular biology grade water, 2x Taqman gene expression master mix (#4369016, Applied Biosystems), and a 20x Taqman gene expression assay probe. The probes used are listed in Table 1. qRT-PCR was performed using the ABI 7300 Real-Time PCR System (Applied Biosystems, CA, USA) and CFX Connect Real-Time System (Bio-Rad Laboratories, CA, USA).

### In-situ hybridization (ISH)

ISH was performed using the RNAscope^®^ HD Reagent Kit-Brown (Advanced Cell Diagnostics, USA). The kit comprised pretreatment reagents, probe detection reagents, wash buffer, and target mRNA probes. The RNAscope^®^ HD assay was conducted according to the manufacturer’s instructions with slight modifications. Specifically, to dehydrate tissue samples, slides were passed through an ethanol gradient of 70%, 85%, 95%, and 100% ethanol. Slides were incubated in an ethanol solution for 5 min. After dehydration, the tissues were cleared by incubating the slides in xylene for 5 min. The slides were mounted in a mounting medium. The results were documented as photomicrographs obtained using a Leica microscope.

### Western blotting (WB)

WB assays were performed using three different protein extracts: 1) total protein, 2) cytoplasmic protein, and 3) nuclear protein. Total protein extracts were prepared using the T-PER™ Tissue Protein Extraction Reagent (#78510, Thermo Fisher Scientific, USA). Cytoplasmic and nuclear protein extracts were prepared using the NE-PER™ Nuclear and Cytoplasmic Extraction Reagent kit (#78833, Thermo Fisher Scientific). The protocol provided by the manufacturer was followed. Protease and phosphatase activities were inhibited by adding the Halt™ Protease and Phosphatase Inhibitor Cocktail (100x) (#78442, Thermo Fisher Scientific) at a final concentration of 1× to the tissue homogenization reagent or buffer. Protein concentrations were measured using the Pierce™ 660 nm protein assay (#22660, Thermo Fisher Scientific). Protein loading samples were prepared using molecular biology grade water, 4x LDS sample buffer (#NP0007, Thermo Fischer Scientific), β-mercaptoethanol, and the respective sample protein extract. The protein concentration of loading samples made from total and cytoplasmic protein extracts was 0.67μg/μL, whereas for those made from nuclear extracts was 0.16 μg/μL. SDS-PAGE was performed using NuPAGE^™^ 4 to 12% Bis-Tris Protein (#NP0336BOX, Thermo Fisher Scientific) pre-cast gels. Following electrophoresis, the separated proteins were transferred onto polyvinylidene fluoride membranes. Membranes were blocked with either 5% milk or BSA in 1xTBST, depending on the primary antibody. Details of the primary antibodies used are presented in Table 2. All secondary antibodies used were conjugated with horseradish peroxidase (HRP); the details are shown in Table 3. Immune complexes were detected using the SuperSignal West Pico PLUS Chemiluminescent Substrate (#34577, Thermo Fisher Scientific). Signals were detected using ImageQuant LAS 4000 Mini (General Electric Life Sciences) and quantified using ImageJ software.

### Immunohistochemistry (IHC)

IHC was performed using a standard routine staining protocol. Briefly, the tissue sections were deparaffinized and rehydrated using xylene and a series of alcohol solutions with decreasing concentration gradients (100%, 95%, 85%, and 70% alcohol). The antigenicity of the target proteins was improved or enhanced by heat-induced epitope retrieval. Tissue sections were boiled in citrate buffer (10 mM citric acid, 0.05% Tween-20, pH 6.0) for approximately 20 min and allowed to cool gradually. Excess peroxidase activity was quenched using 0.3% hydrogen peroxide (H_2_O_2_). The quenching step was skipped for immunofluorescence staining, which used a fluorophore-tagged secondary antibody (2°Ab). Following the quenching step, tissue sections were blocked with an appropriate 5% serum solution in 1X Dulbecco’s phosphate-buffered saline (1X DPBS) with calcium (Ca^2+^) and magnesium (Mg^2+^) (#114-059-101, Quality Biological). Subsequently, the tissue sections were incubated with the appropriate primary antibody (1°Ab) overnight at 4°C. The following day, excess 1°Ab was washed with 1x phosphate buffered saline (1x PBS), and tissues were treated with the appropriate 2°Ab. Detailed descriptions of the 1°- and 2°-Abs used are mentioned in Table 3. For biotin-conjugated 2°Abs, following incubation with 2°Ab, tissue sections were treated with avidin-horse radish peroxidase (HRP) conjugated solution and 3,3’-Diaminobenzamide (DAB) substrate before being counterstained, dehydrated, and mounted. For fluorophore-conjugated 2°Ab, tissue sections were mounted directly using Prolong Gold mounting media containing DAPI counterstain (#P36962, Invitrogen) following 2°Ab treatment.

### Serum collection and blood chemistry analysis

Mice were anesthetized, and blood was extracted from the posterior vena cava. Blood samples were allowed to stand and coagulate for 30 min at room temperature, after which serum was separated by centrifugation. Serum samples were sent to Eli Lilly and Company (Indianapolis, IN, USA) for blood chemistry analysis.

### Statistical analysis

We used the ‘Anova’ test and the ‘Unpaired Student t-test’ (assuming unequal variance) to calculate the p-value and to test for significance. The data obtained were considered statistically significant when compared with the control group (p < 0.05). The error bars in the bar graphs represent standard errors of the mean (SEM).

## Abbreviations

CD133: cell differentiation factor 133
AFP: alpha-fetal protein
EpCAM: epithelial cell adhesion molecules
YAP1: yes-associated protein 1

